# Apelin improves angiogenesis and blood flow reperfusion following lower limb ischemia in diabetic mice

**DOI:** 10.1101/2023.02.08.527688

**Authors:** Stéphanie Robillard, Kien Trân, Tristan Brazeau, Elizabeth Boisvert, Farah Lizotte, Mannix Auger-Messier, Pierre-Luc Boudreault, Éric Marsault, Pedro Geraldes

**Affiliations:** Research Center of the Centre Hospitalier Universitaire de Sherbrooke, Université de Sherbrooke, Québec, Canada; Department of Pharmacology and Physiology, Université de Sherbrooke, Québec, Canada; Division of Cardiology, Université de Sherbrooke, Québec, Canada; Endocrinology, Department of Medicine, Université de Sherbrooke, Québec, Canada

**Keywords:** Peripheral artery disease, angiogenesis, diabetes, apelinergic system, apelin

## Abstract

**BACKGROUND:** Peripheral artery disease (PAD) is a major risk factor for lower-extremity amputation in diabetic patients caused by an insufficient angiogenic response. Unfortunately, therapeutic angiogenesis using growth factors, such as the vascular endothelial growth factor (VEGF), are ineffective in diabetic conditions due to diabetes-induced growth factor resistance. The apelinergic system (APJ receptor/apelin) is highly upregulated under hypoxic condition and acts as an activator of angiogenesis. Apelin treatment has been shown to improve revascularization in nondiabetic models of ischemia, however, its role on angiogenesis in diabetic conditions remains poorly investigated. Thus, this study explored the impact of Pyr-apelin-13 in endothelial cell function and diabetic mouse model of hindlimb ischemia.

**METHODS:** Nondiabetic and diabetic mice underwent femoral artery ligation to induce lower limb ischemia. A group of diabetic mice was implanted subcutaneously with osmotic pumps delivering Pyr-apelin-13 for 28 days. Blood flow reperfusion was measured for 4 weeks post-surgery and exercise willingness was assessed in individual cages with voluntary wheels. *In vitro*, BAECs were exposed to normal (NG) or high glucose (HG) levels and hypoxia. Cell migration, proliferation and tube formation assays were performed following either VEGF or Pyr-apelin-13 stimulation.

**RESULTS:** Following limb ischemia, blood flow reperfusion, functional recovery of the limb and vascular density were improved in diabetic mice receiving Pyr-apelin-13 compared to untreated diabetic mice. In cultured BAECs, exposure to HG concentrations and hypoxia reduced VEGF proangiogenic actions, whereas apelin proangiogenic effects remained unaltered. Pyr-apelin-13 induced its proangiogenic actions through Akt/AMPK/eNOS and RhoA/ROCK signaling pathways under both NG or HG concentrations and hypoxia exposure.

**CONCLUSIONS:** Pyr-apelin-13 promoted endothelial cell function and angiogenesis in the ischemic limb despite diabetes and HG level exposure. Therefore, our results identified the apelinergic system as a potential therapeutic target for angiogenic therapy in diabetic patients with PAD.

**Highlights:**

- Sustained delivery of Pyr-apelin-13 improves blood flow reperfusion, functional recovery of the hindlimb and capillary density in diabetic mice following ischemia.
- Unlike VEGF, signaling pathways and proangiogenic actions induced by apelin/APJ stimulation are not impaired by high glucose exposure.
- Despite the presence of tissue ischemia, diabetes reduces the expression of apelin and APJ in the ischemic adductor muscle, an effect overcome by apelin administration.

## Introduction

Peripheral artery disease (PAD) is defined as a complete or partial atherosclerotic occlusion in the artery, which reduces blood supply to the limbs. As the disease progresses, critical limb ischemia (CLI) and ischemic ulceration of the foot can occur, which increase the risk of limb amputation. Diabetes and smoking are the strongest risk factors for the development of PAD^1^. In individuals with diabetes, PAD appears at an earlier age, progresses more rapidly, and is more severe, which makes this population 5 times more susceptible to lower extremity amputation^2^. Indeed, the combination of PAD and diabetes is responsible for 54% of nontraumatic amputations^3^ and the 5-year survival rate following amputation is 40%^4^. These poor clinical outcomes are partly due to altered angiogenic processes impairing collateral vessel formation, a compensatory mechanism in response to tissue hypoxia. Diabetes causes an imbalance in the production of reactive oxygen species (ROS)^5^, nitric oxide (NO)^6^ and growth factors such as the vascular endothelial growth factor (VEGF)^7^ and the platelet-derived growth factor (PDGF)^8^, contributing to endothelial cell dysfunction. VEGF has been the most studied growth factor for therapeutic angiogenesis and revealed promising results in preclinical studies. However, randomized phase II clinical trials using VEGF gene therapy failed to reduce amputation rate and had no benefits on quality-of-life measurements^9^. These disappointing outcomes could be attributed to diabetes-induced growth factor resistance, inhibiting proangiogenic signaling in endothelial and smooth muscle cells^10^. Therefore, it is crucial to pursue preclinical research for new targets that are not affected by the hyperglycemic milieu to ensure proper collateral vessel formation and reduce the incidence of CLI and limb amputation.

Apelin and its receptor APJ, a G protein-coupled receptor (GPCR), are expressed in multiple tissues such as the heart, brain, kidney, adipose tissue and endothelium^11^. Being widely distributed in the body, apelin/APJ modulate multiple physiological processes including cardiac inotropy, blood pressure, glucose metabolism, water homeostasis and angiogenesis^12^. The main apelin fragments generated by the cleavage of preproapelin are apelin-13, apelin-17 and apelin-36, and they all differently regulate APJ signal transduction pathways, causing different biological effects^13,14^. Apelin-13 can be further post-translationally modified by a cyclization of the glutamine in N-terminus to produce pyroglutamylated apelin-13 (Pyr-apelin-13)^15^. Compared to the other apelins, Pyr-apelin-13 is the most potent^16^ and abundant isoform in the plasma and cardiac tissue^16,17^ with enhanced plasma half-life^18^. Apelin/APJ axis is essential for vascular homeostasis since it regulates the enlargement of new blood vessels during angiogenesis and regulates parallel alignment of arteries and veins in the skin^19^. Furthermore, APJ has been shown to be required for normal development of the cardiovascular system since in APJ deficient mice, over fifty percent of embryos die during pregnancy and the survived embryos have insufficient vascular maturation and abnormal ventricular wall formation^20^.

Apelin administration or apelin gene therapy has been reported to improve revascularization in nondiabetic model of hindlimb ischemia^21-23^. However, to our knowledge, no study has investigated the impact of the apelinergic system (apelin/APJ axis) on angiogenesis in a diabetic PAD model. The present study further explored the contribution of Pyr-apelin-13 on angiogenesis in endothelial cells exposed to high glucose concentrations and hypoxia, and in diabetic mice following hindlimb ischemia.

## Research Design and Methods

### Reagents and antibodies

Primary antibodies for immunoblotting were purchased from commercial sources: GAPDH horseradish peroxidase (V-18), endothelial nitric oxide synthase (eNOS) (C-20) from Santa Cruz Biotechnology Inc (Dallas, TX); protein kinase B (Akt) (9272S), phospho-Akt (193H12), AMP-activated protein kinase *α* (AMPK*α*) (2532S), p-AMPK*α* (2535S), p-eNOS Ser1177 (9571S), Ras homolog family member A (RhoA) (2117S), rho-associated coiled-coil-containing protein kinase (ROCK) 1 (4035S), ROCK-2 (8236S), and secondary antibody of anti-rabbit and anti-mouse peroxidase-conjugated from Cell Signaling (Danver, MA); anti-α smooth muscle actin (ab5694) and p-ROCK2 (ab228008) from Abcam (Toronto, ON); anti-CD31(558736) and p-eNOS Ser633 (612664) from BD Bioscience (Mississauga, ON). Secondary antibodies for immunofluorescence FITC conjugated anti-rat IgG and Alexa Fluor 647 conjugated anti-rabbit IgG were purchased from Jackson ImmunoResearch Laboratories (West Grove, PA). Halt Protease Inhibitor Cocktail (78430) was purchased from Thermo Fisher Scientific (Waltham, MA). SEP-COLUMN (RK-SEPCOL-1) for plasma peptides extraction and Apelin-12 (Human, Rat, Mouse, Bovine) Enzyme Immunoassay (EIA) (EK-057-23) were purchased from Phoenix Pharmaceuticals Inc (Burlingame, CA). Fetal bovine serum (FBS), penicillin-streptomycin (P/S), phosphate-buffered saline (PBS), and Dulbecco’s Modified Eagle Medium (DMEM) low glucose (31600-034) were obtained from Invitrogen (Burlington, ON). VEGF-A_165_ was purchased from R&D (Minneapolis, MN). Pyr-apelin-13 (pyr-Ape-13) was synthesized and generously provided by Dr. Boudreault’s laboratory from the *Institut de Pharmacologie de Sherbrooke*. All other reagents used, including streptozotocin, ethylenediaminetetraacetic acid (EDTA), bovine serum albumin (BSA) d-glucose, d-mannitol, leupeptin, phenylmethylsulfonyl fluoride, aprotinin, and Na_3_VO_4_ were purchased from Sigma-Aldrich (St. Louis, MO).

### Cell culture

Bovine aortic endothelial cells (BAECs) were isolated from fresh harvested aorta as previously described^7^. Cells from passages 2 to 7 were trypsinized and cultured in DMEM 2.5% FBS and 1% penicillin-streptomycin. For all the *in vitro* experiments, cells were cultured in DMEM 0.1% FBS containing normal (NG; 5.6 mmol/L) or high glucose (HG; 25 mmol/L) levels for up to 48 hours and exposed to hypoxia for the last 16 hours (1% O_2_) to reproduce the ischemic state of PAD. BAECs were stimulated with either VEGF-A (10 ng/mL) for 5 minutes or Pyr-apelin-13 (200 nM) for 1h (immunoblotting and cell signaling experiments) or 24h prior to RNA extraction and quantitative PCR analysis.

### Migration assay

BAECs were trypsinized, counted and seeded at 20 000 cells inside the 2 wells of Ibidi’s culture insert (80209, Ibidi) placed into a 12-well plate. After cell adhesion (4 hours later), BAECs were exposed to NG or HG concentrations for 24h. Following the 24h treatment, each culture insert was removed, and the wells were filled with 1 ml of either NG or HG medium. Cells were then stimulated, or not, with either VEGF-A (25 ng/mL) or Pyr-apelin-13 (100 nM) and placed into the hypoxic incubator (1% O_2_) for 16h. Cell migration was evaluated by taking images under Nikon eclipse Ti microscope at 10X magnification immediately after removing the culture insert and at the end of the experiment (16h later). Analysis was performed with Image J software by measuring the difference in occupied area immediately after insert removal and following 16h stimulation in NG or HG condition. Results were reported as a % of cell migration for analysis.

### Proliferation assay

BAECs were trypsinized, counted and seeded in a 96-well plate at 5 000 cells per well. After cell adhesion (4 hours later), cells were exposed to NG or HG concentrations for 24h, stimulated with either VEGF-A (25 ng/mL) or Pyr-apelin-13 (100 nM) and placed into the hypoxic incubator for 16h. Cells were then fixed in 4% paraformaldehyde for 5 min, rinsed twice in PBS and incubated with the nuclear counterstain DAPI (Sigma-Aldrich) at 0.001 mg/ml for 10 min. Fluorescence microscopy and the NIS-Elements software of Nikon eclipse Ti microscope were used to visualize and count cells, reported as the number of cells/mm^2^ for analysis.

### Tube formation assay

BAECs were cultured in 100 mm petri dish and exposed to NG or HG concentrations for 24h and then placed into the hypoxic incubator (1% O_2_) for 16h. On the day of the assay, 10 *μ*l of Matrigel Matrix (Growth Factor Reduced, 356230, Corning) was applied into each well of Ibidi’s *μ*-slide rack (81506, Ibidi) and incubated for 30 minutes at 37°C to allow polymerization. Cells were then trypsinized, counted, and 600 000 cells were seeded per well on top of the Matrigel. After seeding, cells were immediately stimulated, or not, with VEGF-A (25 ng/mL) or Pyr-apelin-13 (100 nM) and the *μ*-slide rack was placed into the hypoxic incubator for 4h. At the end of the experiment (4h later), the tube formation was visualized, and images were taken under Nikon eclipse Ti microscope at 4X magnification. Endothelial cells tube formation ability after NG or HG treatment and angiogenic factor stimulation was measured using Image J software by counting the number of formed tubes and normalized on the NG condition to present it as a fold increase.

### Immunoblot analysis

Adductor muscles or endothelial cells were lysed in RIPA buffer containing protease inhibitors (1 mmol/l phenylmethylsulfonyl fluoride, 2 µg/ml aprotinin, 10 µg/ml leupeptin, 1 mmol/l Na3VO4, 1 mmol/l NaF) and protein concentrations were measured by DC kit (BioRad). The lysates (20 *μ*g for BAECs and 50 *μ*g for adductor muscles) were separated by SDS-PAGE and transferred to PVDF membranes, which were blocked with 5% skim milk for 1 hour at room temperature. Membranes were incubated overnight with primary antibodies at a 1:1000 dilution in 5% skim milk (or in 5% BSA for phospho-ROCK-2 and p-AMPK*α*), followed by the corresponding secondary antibodies conjugated with horseradish peroxidase at a 1:10 000 dilution in 5% skim milk (or 1:5 000 for phospho-ROCK-2 and p-AMPK*α*). Immobilon Forte Western HRP substrate (millipore, Etobicoke, ON) was used to visualize immunoreactive bands and protein content was quantified using Computer-assisted densitometry ImageLab imaging software (Chemidoc, BioRad).

### Animal and experimental design

C57Bl/6 male mice were purchased from Charles River (strain 027). At 8 weeks of age, the mice were rendered diabetic by intraperitoneal streptozotocin injection (50 mg/kg in 0.05 mol/L citrate buffer, pH 4.5; Sigma) on 5 consecutive days after overnight fasting. Control mice were injected with citrate buffer. Blood glucose was measured with a glucometer (Contour, Bayer, Inc.) one week after the injections to confirm diabetes. All experiments were conducted in accordance with the Canadian Council of Animal Care and were approved by the Animal care and Use Committees of the Université de Sherbrooke, according to the NIH Guide for the Care and Use of Laboratory Animals.

### Hindlimb ischemia model

After 2 months of diabetes, blood flow was measured in 16-week-old nondiabetic (NDM) and diabetic (DM) mice. Animals were anesthetized by inhalation of Isoflurane USP (1-chloro-2,2,2-trifluoroethyl difluoromethyl ether) at a concentration of 5% (initiation) and then maintained at 1-2% during the whole surgical procedure (approximately 20 minutes). To mimic the ischemic condition in PAD, unilateral hindlimb ischemia was induced by ligation of the femoral artery as we previously described^24^. Directly after the hindlimb surgery, osmotic pumps (model 1004; Alzet Osmotic pumps) with Pyr-apelin-13 (1 mg/kg/day in PBS) were implanted subcutaneously, according to the manufacturer’s instructions, to allow 4-week infusions. The blood flow recovery following hindlimb ischemia was evaluated in 3 groups of mice: NDM, DM, DM+pyr-Ape-13.

### Laser doppler perfusion imaging

The hindlimb blood flow was measured using a laser Doppler perfusion imaging (PIMIII) system (Perimed Inc.). Consecutive perfusion measurements were obtained by scanning the region of interest (hindlimb and foot) of anesthetized animals. Measurements were performed pre-artery and post-artery ligation to ensure the success of the surgery, and on post-operative days 7, 14, 21 and 28. To account for variables that affect blood flow temporally, the results at any given time were expressed as a ratio against simultaneously obtained perfusion value of the ischemic (right) and nonischemic (left) hindlimb. Following laser Doppler perfusion imaging on day 28, mice were euthanized by exsanguination via the left ventricle under deep anesthesia (Isoflurane USP, inhalation at a concentration of 5%) and the ischemic adductor muscles were harvested.

### Voluntary exercise wheel

Mice were housed individually for 5 consecutive days between day 23 and 27 to measure their activity using voluntary exercise wheel (Scurry mouse activity wheel, Model 80820FS, Lafayette Instrument Neuroscience). A magnetic sensor (Scurry mouse activity counter, Model 86110) attached to the wheel recorded the number of revolutions and running distance using the Scurry activity monitoring software (Lafayette Instrument Neuroscience). Results are shown as cumulative distance of the 5 days.

### Apelin peptide levels in the plasma

Mice were euthanized by exsanguination and blood was immediately centrifuged to isolate the plasma. Halt Protease Inhibitor Cocktail was added to the plasma to prevent apelin degradation and samples were snap freeze in liquid nitrogen. Peptides from the plasma were extracted on SEP-COLUMN containing 200 mg of C18 following the manufacturer’s instructions. Briefly, the plasma was acidified using the Buffer A provided in the extraction kit and loaded into the pretreated C-18 SEP-COLUMNs. Columns were washed twice with Buffer A then the peptides were eluted with Buffer B. Eluants were evaporated to dryness in a centrifugal concentrator and then rehydrated in 125 ul of Assay Buffer provided in the EIA kit. Apelin-12 EIA kit was used according to the manufacturer’s procedures to detect apelin-12, 13 and 36 isoforms, with a sensitivity level in the range of ng/mL, based on the principle of competitive EIA.

### Histopathology

Ischemic adductor muscles from NDM, DM and DM+pyr-Ape-13 were harvested for pathological examination and sections were fixed in 4% paraformaldehyde (VWR) for 18 h and then transferred to 70% ethanol. Fixed tissues were embedded in paraffin and 4 µm sections were stained with hematoxylin & eosin (H&E) or used for immunofluorescence assay.

### Immunofluorescence

Cross-sections of ischemic adductor muscle were blocked at room temperature for an hour with 10% goat serum and then incubated overnight with primary antibodies (CD31 (1:50) and α-smooth muscle actin, 1:200), followed by 1h incubation with the secondary antibody (1:400). Images of the entire ischemic muscle were taken with the Nikon eclipse Ti microscope and the number of vessels smaller than 30 *μ*m in lumen diameter was counted on each image. All images were taken at the same time under identical settings and similarly handled in Adobe Photoshop and Image J software.

### Quantitative PCR analysis

Quantitative PCR was performed to evaluate mRNA expressions of genes in the ischemic adductor muscles and BAECs as we previously described^7^. PCR primers are listed in Supplemental Table 1. The expression of the housekeeping gene glyceraldehyde 3-phosphate dehydrogenase (GAPDH) was used to normalized data.

### Statistical analyses

*In vivo* and *in vitro* data are shown as mean ± SD for each group except for blood flow measurements which are presented as mean ± SEM. Statistical analysis was performed by unpaired Kruskal-Wallis followed by Dunn’s multiple comparisons test (Figure 1B, 1C, 1E and 1F, Figure 3A through D, Figure 4A and D and Figure 5G) or by unpaired One-Way ANOVA followed by Tukey’s multiple comparisons test (Figure 2B through 2D, 2G and 2H, Figure 3E and 3F, Figure 4B, 4C and 4E through 4H, Figure 5A through 5F).

**Figure 1.**
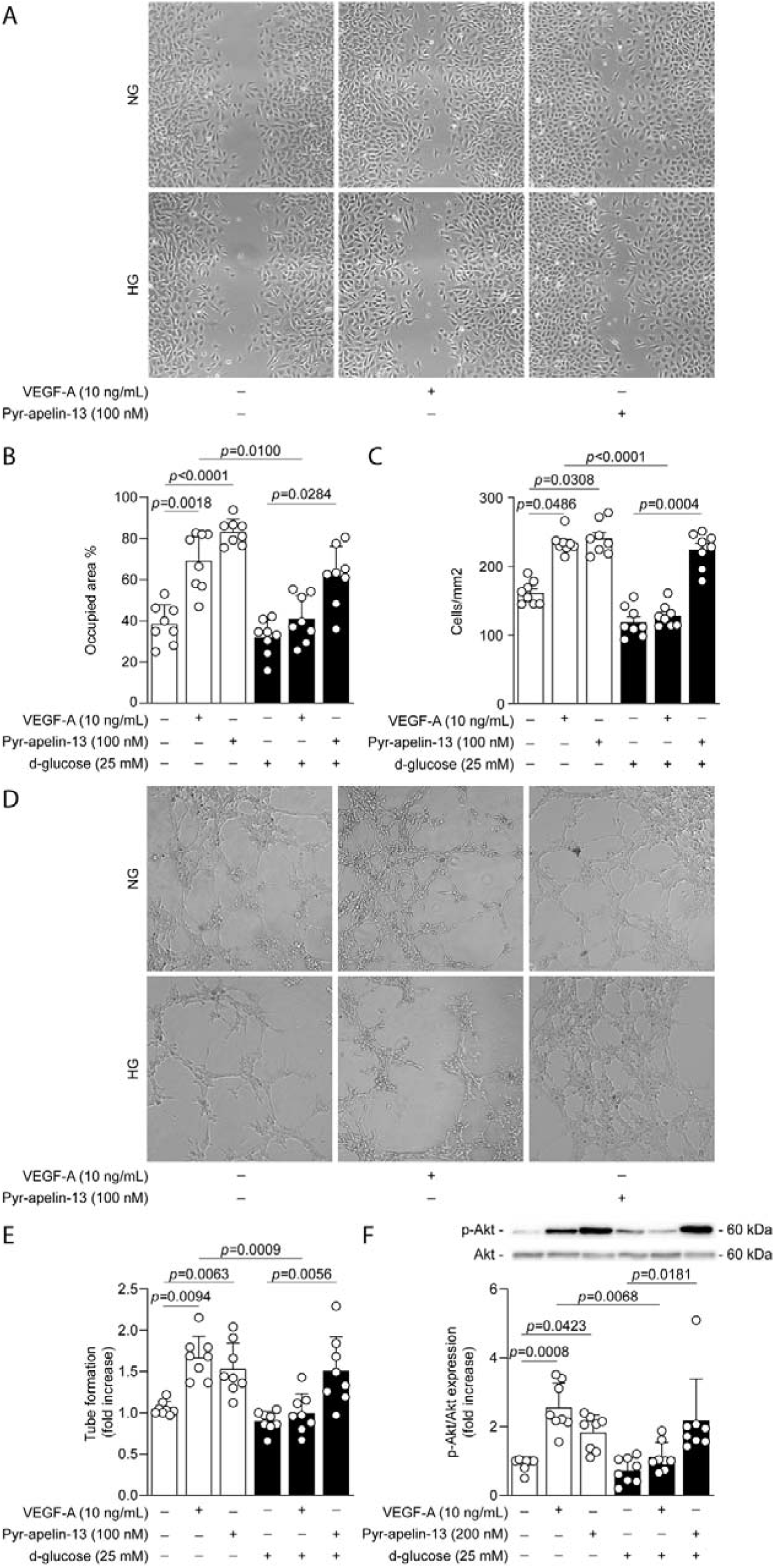
High glucose exposure did not affect Pyr-apelin-13 proangiogenic action in endothelial cells under hypoxia. BAECs were incubated with normal glucose (NG; 5.6 mmol/L; white bars) or high glucose (HG; 25 mmol/L; black bars) for up to 48h and then stimulated with either VEGF-A or Pyr-apelin-13 for 16h **(A-C)**, 4h **(D-E)**, 5 min (VEGF) or 1h (Pyr-apelin-13) **(F)** under hypoxia (1% O_2_) for the last 16h of treatment in all experiments. **(A)** Representative images of the cell migration assay using the Ibidi’s insert. **(B)** The percentage of the surface area occupied by the BAECs was quantified. **(C)** BAECs were fixed and stained with DAPI (4’,6-diamidino-2-phenylindole) and then cells were counted using the NIS-Elements software of Nikon eclipse Ti microscope. **(D)** Representative images of the tubule formation abilities of endothelial cells using Ibidi’s *μ*-slide rack. **(E)** Tubule formation was quantified by measuring the total number of closed circles in the entire well, normalized on the NG condition. **(F)** Protein expression of Akt phosphorylation was detected by immunoblot analysis and the densitometry quantification was measured. Results are presented as the mean ± SD of 8 independent cell experiments.

**Figure 2.**
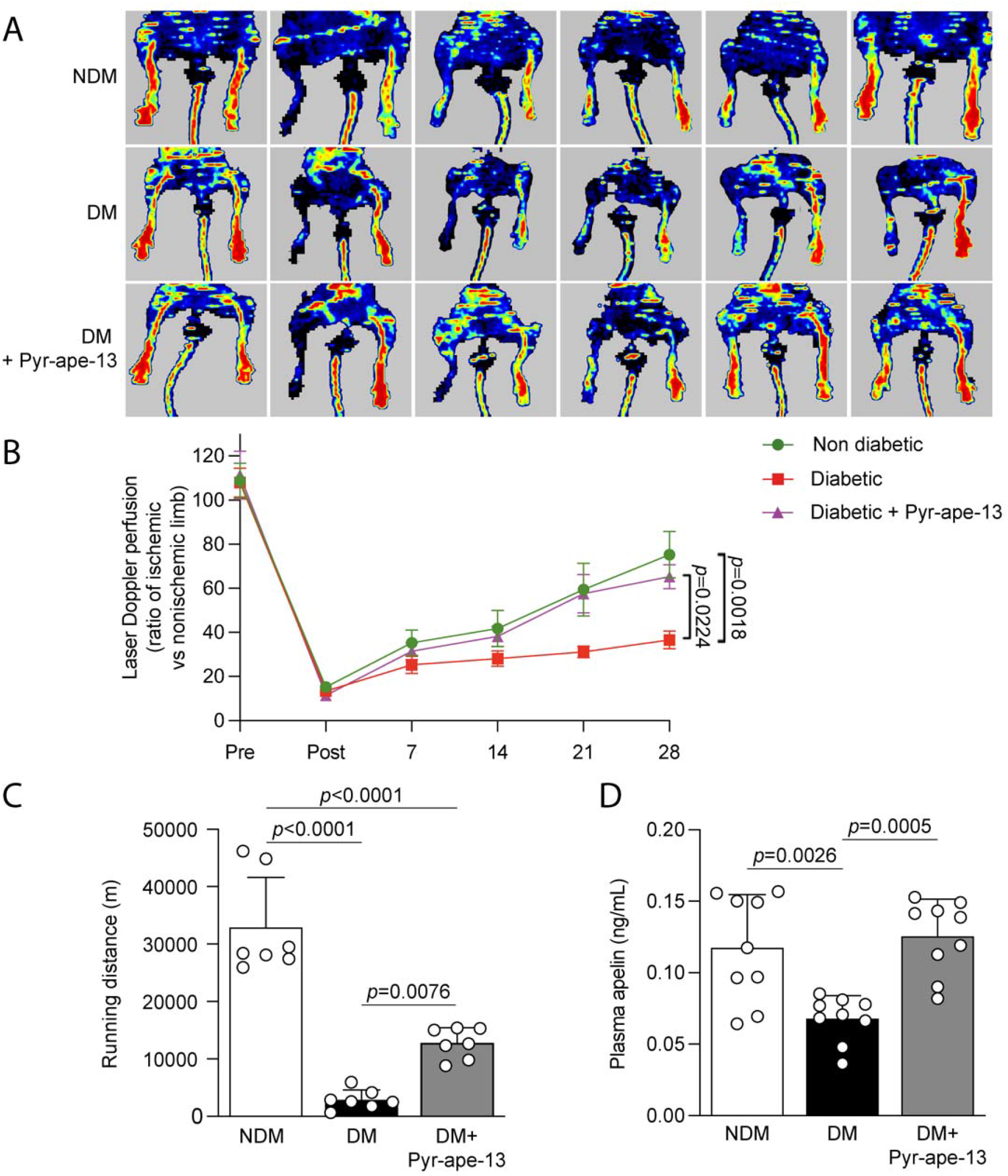
Pyr-apelin-13 delivery resulted in improved blood flow reperfusion, running distance, vascular density and muscle structure in diabetic mice following critical limb ischemia. **(A)** Laser Doppler imaging and **(B)** blood flow reperfusion analysis of nondiabetic (NDM), diabetic (DM) and diabetic mice receiving Pyr-apein-13 (DM+Pyr-ape-13), pre, post, and 4 weeks following femoral artery ligation. **(C)** Cumulative running distance (m) over 5 days in the voluntary exercise wheel made by nondiabetic (NDM; white bars), diabetic (DM; black bars) and diabetic mice receiving Pyr-apelin-13 (DM+Pyr-ape-13; grey bars). (**D)** Apelin plasma levels in ng/mL. Results are presented as the mean ± SEM of 12 mice per group **(B)** and as the mean ± SD of 7 **(C)** and 9 **(D)**.

**Figure 3.**
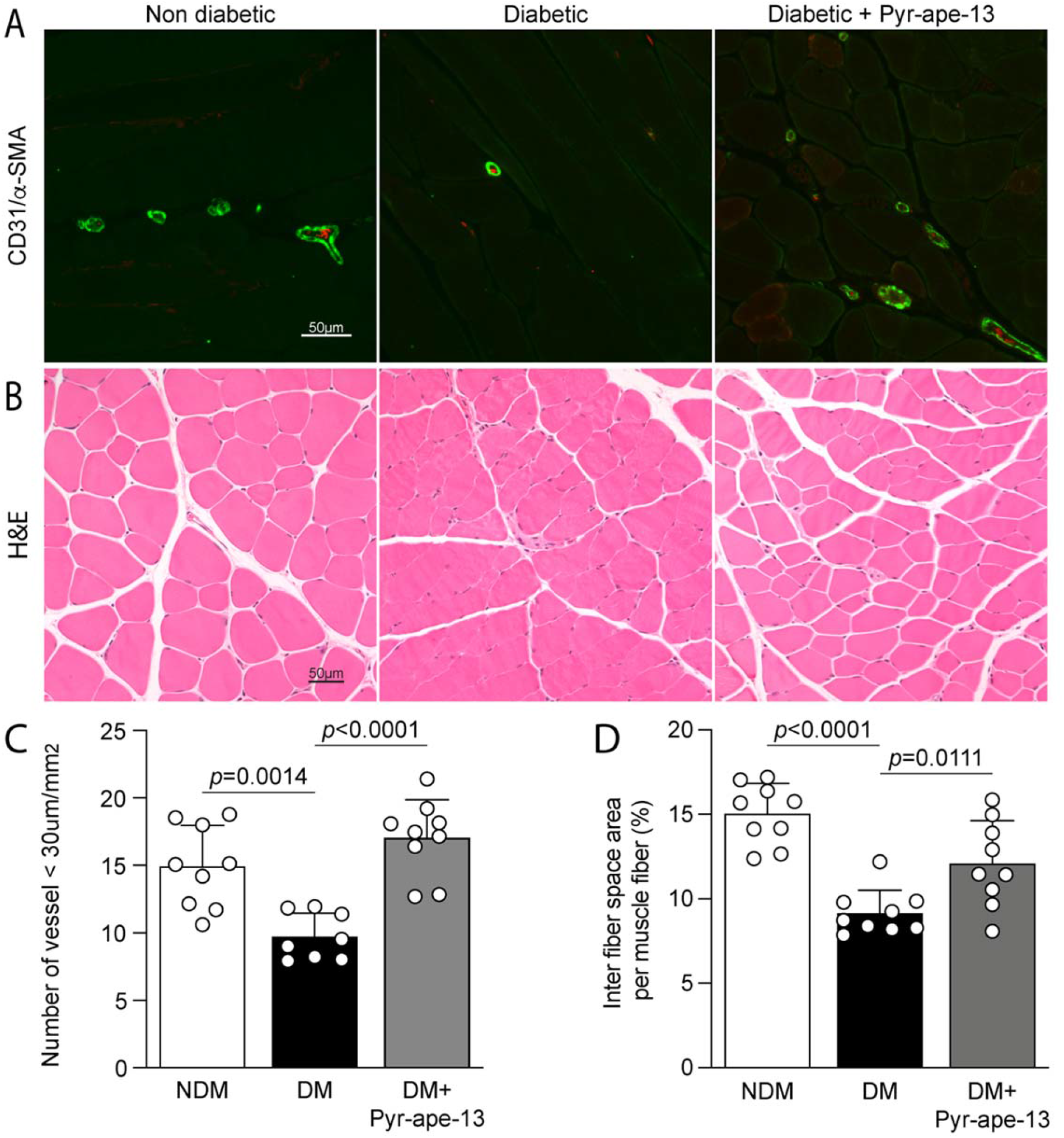
Pyr-apelin-13 administration in diabetic mice enhanced vascular density and improved muscle structure integrity following critical limb ischemia. **(A)** Immunofluorescence images of endothelial cells (CD31; red) and a-smooth muscle actin (green). **(B)** Structural analysis of the ischemic muscles stained with hematoxylin and eosin (H&E). **(C)** quantification of the number of vessels smaller than 30 *μ*m in the ischemic muscle. **(D)** Quantification of the space between muscle fibers of the ischemic muscles. Results are presented as the mean ± SD of 8-9 **(A** and **C)** and 9 mice per group **(B** and **D)**.

**Figure 4.**
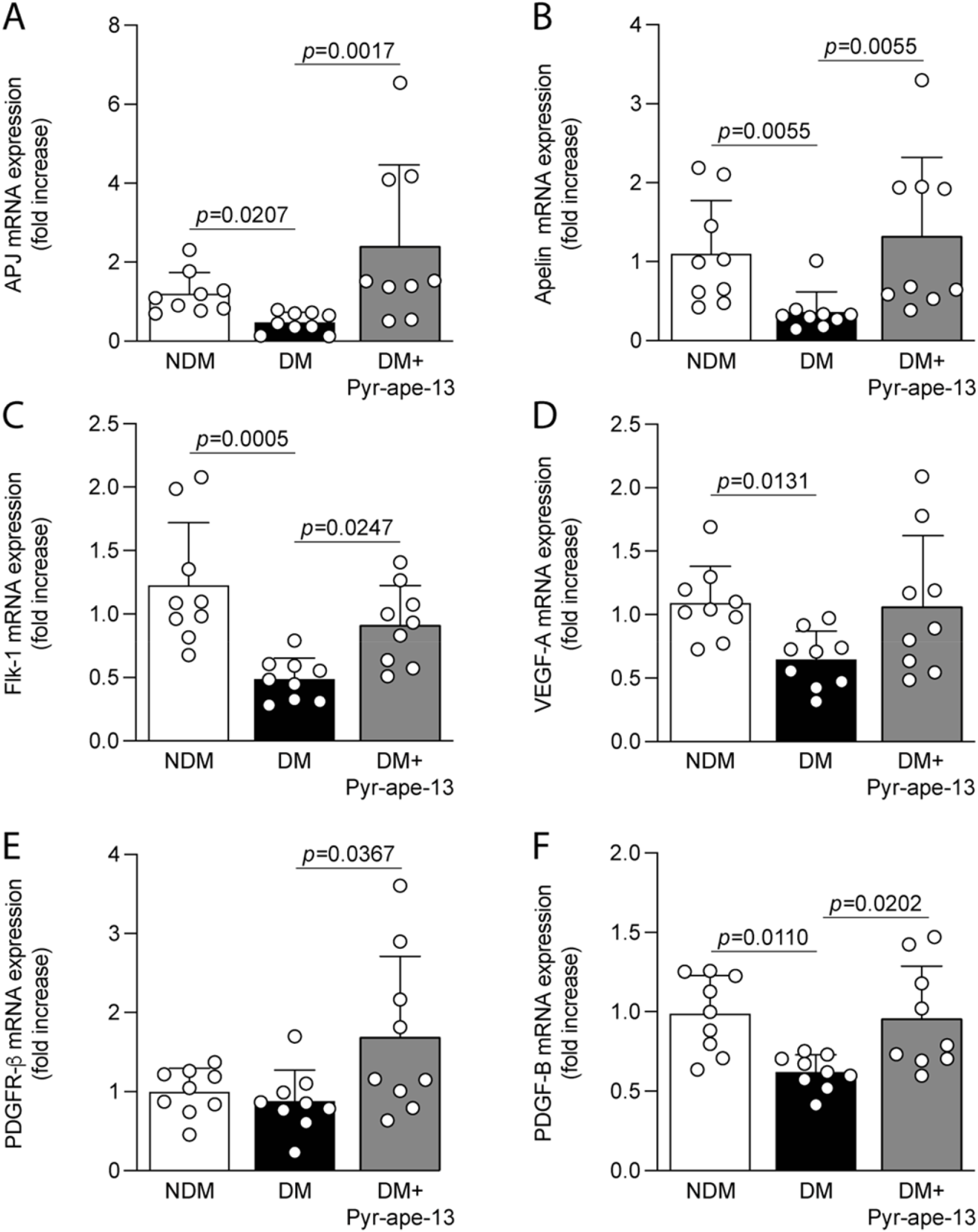
Pyr-apelin-13 increased the expression of growth factors in the ischemic muscle from diabetic mice following ischemia. Expression levels of **(A)** APJ, **(B)** Apelin, **(C)** Flk-1, **(D)** VEGF-A, (**E)** PDGFR-*β* and **(F)** PDGF-B mRNA in the ischemic muscle of nondiabetic (NDM; white bars), diabetic (DM; black bars) and diabetic mice receiving Pyr-apelin-13 (DM+Pyr-ape-13; grey bars). GAPDH gene was used for mRNA normalization. Results are presented as the mean ± SD of 9 mice per group.

**Figure 5.**
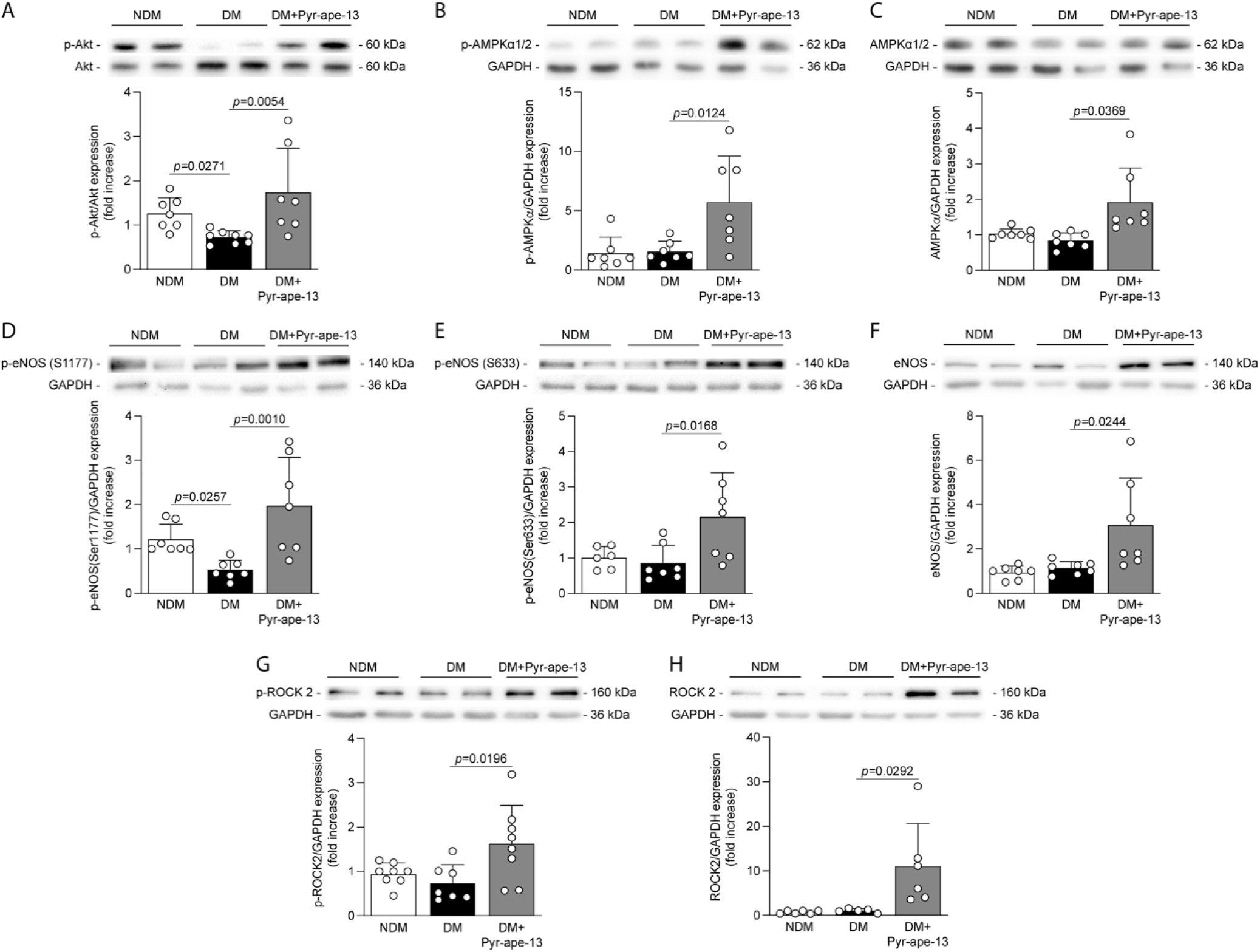
Pyr-apelin-13 delivery in diabetic mice promoted angiogenesis following limb ischemia through the activation of Akt, AMPK, eNOS and ROCK2 signaling pathways. Protein expression of **(A)** phospho-Akt, **(B)** phosphor-AMPK*α*1/2, **(C)** AMPK*α*1/2, **(D)** phospho-eNOS at Ser1177, **(E)** phospho-eNOS at Ser633, **(F)** eNOS, **(G)** phospho-ROCK2 and **(H)** ROCK2 in the ischemic adductor muscle of nondiabetic (NDM; white bars), diabetic (DM; black bars) and diabetic mice receiving Pyr-apelin-13 (DM+Pyr-ape-13; grey bars) was detected by immunoblot analysis and the densitometry quantification was measured. Results are presented as the mean ± SD of 7-8 **(A** and **G)**, 7 **(B-D** and **F)**, 6-7 **(E)** and 5-6 mice per group **(H)**.

## Results

### Proangiogenic properties of Pyr-apelin-13 were not affected by high glucose exposure in endothelial cells

We have previously demonstrated that crucial factors involved in the angiogenic processes, such as VEGF, are affected in diabetes^7,25^. To evaluate and understand if apelin can improve endothelial function under a hyperglycemic milieu, we have evaluated its role on critical steps involved in blood vessel formation and compared its response to VEGF-A. Under NG conditions, hypoxia exposure promoted endothelial cell migration by 38% coverage of the surface area as compared to the surface area initially created by Ibidi’s insert. As expected, under the same conditions, VEGF-A and Pyr-apelin-13 stimulation enhanced cell migration by 69% (*P*=0.0018) and 83% (*P*<0.0001), respectively (Fig. 1A and 1B). As previously reported under HG and hypoxic conditions, the stimulation with VEGF-A only raised cell migration by 41% (*P*>0.9999), while Pyr-apelin-13 promoted endothelial cell migration by 62% (*P*=0.0284; Fig. 1A and 1B). Cell proliferation is also an important step in the angiogenic process. Our data demonstrated that the promitogenic effect of VEGF-A (1.4-fold; *P*=0.0486) on endothelial cell proliferation under NG and hypoxic conditions were blunted in HG level exposure (Fig. 1C). In contrast, Pyr-apelin-13-induced endothelial cell proliferation was conserved in HG conditions (1.9-fold; *P*=0.004; Fig. 1C). Then, we investigated the capacity of these growth factors to induce vascular tubule formation on Matrigel. We observed that VEGF and apelin treatment increased similarly endothelial cell tubule formation when exposed to NG condition and hypoxia by 1.6-fold (*P*=0.0094) and 1.5-fold (*P*=0.0063), respectively (Fig. 1D and 1E). In line with the other angiogenic processes, the capacity of VEGF to induce tubule formation was abrogated in HG level exposure, which was not observed with Pyr-apelin-13. Treatment with Pyr-apelin-13 enhanced tubule formation in both NG (1.5-fold; *P*=0.0063) and HG (1.7-fold; *P*=0.0056) conditions (Fig. 1D and 1E). Inhibition of VEGF-induced proangiogenic actions in diabetes has been well documented by us and others, and associated with impaired activation of VEGFR-2 and downstream effector Akt^7,26^. As we reiterated in our study, under NG and hypoxic conditions, VEGF stimulation led to a 2.8-fold increase (*P*=0.0008) in Akt phosphorylation, whereas under HG exposure, Akt activation was reduced by 56% (*P*=0.0068; Fig. 1F). In contrast to VEGF-A, Pyr-apelin-13 stimulated Akt phosphorylation by 2-fold (*P*=0.0423) and 3-fold (*P*=0.0181) under hypoxia and NG or HG conditions, respectively (Fig.1F). Taking together, these results clearly demonstrate that HG conditions caused inhibition of VEGF actions on endothelial cell proangiogenic function, a phenomenon that was not observed under the same conditions following Pyr-apelin-13 stimulation.

### Pyr-apelin-13 enhanced blood flow reperfusion following hindlimb ischemia in diabetic mice

Since *in vitro* experiences in endothelial cells exposed to HG levels indicated that treatment with apelin remained effective to induce proangiogenic actions, we investigated its potential beneficial effects in a diabetic hindlimb ischemia mouse model. Treatment with apelin osmotic pumps did not influence systemic blood glucose nor affect weight in diabetic mice (Supplemental Table 2). After 2 months of diabetes, femoral artery ligation was performed in nondiabetic and diabetic mice receiving or not systemic delivery of Pyr-apelin-13 with subcutaneous osmotic pumps. Blood flow reperfusion of the lower limb was measured once a week over a period of 4 weeks using laser Doppler imaging (Fig. 2A). Following 28 days post-ligation, diabetic mice exhibited a 36% blood flow recovery compared to 75% in nondiabetic mice (*P*=0.0018; Fig. 2B). Interestingly, systemic administration of Pyr-apelin-13 in diabetic animals improved blood flow reperfusion to 65% (*P*=0.0224; Fig. 2B).

### Pyr-apelin-13 improved motor function of the ischemic hindlimb in diabetic mice

A common symptom in patients with PAD is the development of fatigue or pain in the legs leading to reduced walking distance. Thus, we have evaluated the functional recovery by placing the mice in individual cages equipped with a voluntary exercise wheel, an environment without constraint and human intervention, during 5 consecutive days. Three weeks following limb ischemia, untreated diabetic mice exhibited a significant 90% reduction (*P*<0.0001) in running distance as compared to nondiabetic mice (Fig. 2C). Interestingly, diabetic mice receiving Pyr-apelin-13 improved by 4.4-fold (*P*=0.0076) compared to untreated diabetic mice (Fig. 2C). These results suggested an impact of diabetes on the functional recovery of the hindlimb following ischemia, that can be partially recovered by the administration of Pyr-apelin-13 in diabetic mice.

### Diabetes reduced plasma apelin levels following ischemia

Apelin levels in the plasma were measured using Phoenix Pharmaceutical EIA kit, reported to cross-react with apelin-12, 13 and 36 isoforms. Several studies reported variations in apelin plasma concentrations in the context of diabetes^27,28^. We observed a 42% reduction (*P*=0.0026) in apelin plasma levels in diabetic mice in response to limb ischemia compared to nondiabetic mice (Fig. 2D). Implantation of Alzet osmotic pumps immediately after limb surgery ensured a continuous systemic delivery of apelin for a period of 28 days, which enhanced plasma apelin concentrations in diabetic mice by 1.8-fold (*P*<0.0001) compared to untreated diabetic mice (Fig. 2D).

### Pyr-apelin-13 increased vascular density in the ischemic adductor muscle of diabetic mice

To support the blood flow reperfusion data, we evaluated the ability of Pyr-apelin-13 to promote collateral vessel formation following ischemia by measuring vascular density on cross-sections of the ischemic adductor muscle of all groups of mice. Diabetes reduced the formation of blood vessels (< 30 *μ*m) in response to tissue ischemia by 35% (*P*=0.0014) compared to normoglycemic littermate controls (Fig. 3A and 3C). However, delivery of Pyr-apelin-13 in diabetic mice significantly enhanced the amount of blood vessels, from 9.74 to 17.05 vessels/mm^2^ (1.8-fold, *P*<0.0001; Fig. 3A and C), suggesting that Pyr-apelin-13 treatment can promote angiogenesis in response to ischemia despite being exposed to diabetes. In addition, we measured the interfiber space area on H&E staining sections of the ischemic muscle as an indication of general muscle structure. We observed differences in the structural integrity of the muscle fibers from diabetic mice (P<0.0001) compared to nondiabetic mice (Fig. 3B and 3D). Interestingly, apelin delivery improved muscle structural integrity and fiber organization by increasing interfiber space area in diabetic mice by 1.3-fold (*P*=0.0111; Fig. 3B and 3D).

### Treatment with Pyr-apelin-13 enhanced growth factor expression in diabetic ischemic hindlimb

Apelin and its receptor are known to be upregulated in the muscle and the artery by hypoxia in nondiabetic animal models^21,22^. Gene expression of the apelinergic system is influenced by diabetes in different tissues^27,29^. Following 28 days of femoral artery ligation, we observed that diabetes reduced mRNA expression of both APJ and apelin in the ischemic muscle by 60% (*P*=0.0207; Fig. 4A) and 67% (*P*=0.0055; Fig. 4B), respectively. In contrast, Pyr-apelin-13 administration increased APJ mRNA expression by 3.4-fold (*P*=0.0017; Fig. 4A) and its own expression by 3.6-fold (*P*=0.0055; Fig. 4B) in diabetic mice. As we previously reported, many other growth factors are downregulated by diabetes in the ischemic muscle, including VEGF-A, Flk-1, PDGF-B and PDGFR-*β*^7,8,24^. Increased capillary density in the ischemic hindlimb of diabetic mice receiving Pyr-apelin-13 led us to investigate whether apelin influenced the expression pattern of well-known proangiogenic growth factors. In our study, Flk-1 (*P*=0.0005; Fig. 4C), VEGF-A (*P*=0.0131; Fig. 4D) and PDGF-B (*P*=0.0110; Fig. 4F) mRNA expression in the ischemic muscle were significantly decreased by diabetes as compared to nondiabetic controls. Our results indicated that the administration of Pyr-apelin-13 significantly increased the gene expression of Flk-1 (*P*=0.0247; Fig. 4C), VEGF-A (*P*=0.0746; Fig 4D), PDGFR-*β* (*P*=0.0367; Fig. 4E) and PDGF-B (*P*=0.0202; Fig. 4F) in diabetic mice.

### Pyr-apelin-13 promoted revascularization in vivo through proangiogenic and cytoskeleton organization signaling pathways

The signaling pathways by which apelin may promote angiogenesis under hyperglycemia were investigated in the ischemic adductor muscle. The binding of apelin to its receptor APJ, a GPCR, leads to the recruitment of the small G*α*_i_ protein which subsequently activates the Akt signaling pathway, promoting cell survival and angiogenesis. As expected, diabetes markedly reduced Akt phosphorylation in the ischemic muscle by 42% (*P*=0.0271) compared to nondiabetic mice (Fig 5A). In contrast, diabetic mice who received Pyr-apelin-13 exhibited a 2.4-fold (*P*=0.0054) increase Akt phosphorylation (Fig. 5A). APJ activation may also induce the phosphorylation and activation of AMPK via the G*α*_i_ and G*α*_q_ subunits^30^. AMPK is mainly known as a fuel-sensing enzyme and for its metabolic functions, but it also promotes endothelial cell proliferation and migration^31^. Our results showed that apelin delivery raised AMPK*α*1/2 phosphorylation by 3.7-fold (*P*=0.0124; Fig. 5B) and total protein expression by 2.2-fold (P=0.0369; Fig. 5C) in the ischemic muscle of diabetic mice compared to untreated diabetic mice. Then, we evaluated eNOS as a common downstream target of Akt and AMPK. Once phosphorylated, eNOS produces NO, a major regulator of endothelial cell growth and angiogenesis^32^. It has been shown that Ser 633 and Ser1177 of eNOS can be phosphorylated by AMPK while Akt only phosphorylates eNOS at Ser1177^33,34^. As expected, diabetes reduced Ser1177 phosphorylation of eNOS by 60% (*P*=0.0257; Fig. 5D) compared to normoglycemic animals but had little impact on eNOS Ser633 activation. Interestingly, the administration of Pyr-apelin-13 in diabetic mice increased both Ser1177 and Ser633 phosphorylation of eNOS by 4.6-fold (*P*=0.0010; Fig. 5D) and 2.5-fold (*P*=0.0168; Fig. 5E), respectively. Furthermore, apelin delivery in diabetic mice elevated total eNOS protein expression in the ischemic muscle by 2.7-fold (*P*=0.0244) compared to untreated diabetic mice (Fig. 5F). We also explored the RhoA/ROCK signaling pathway, which is known to be recruited by the small G*α*_12/13_ protein following GPCRs activation^35^, and can mediate cellular morphological processes such as cell contractility, actin cytoskeleton organization, cell migration and proliferation^36^. Although no difference in ROCK-2 phosphorylation and protein expression was observed in the ischemic muscle of diabetic mice compared to nondiabetic mice, apelin administration raised ROCK2 phosphorylation by 2.2-fold (*P*=0.0196; Fig. 5G) and ROCK-2 total protein expression by 10.7-fold *(P*=0.0292; Fig. 5H) compared to untreated diabetic mice.

### Pyr-apelin-13 enhanced cultured endothelial cells proangiogenic functions through the Akt/AMPK/eNOS and RhoA/ROCK2 pathways

To confirm that the signaling pathways enhanced by the treatment with Pyr-apelin-13 in ischemic muscle occurred in endothelial cells, we investigated their activation in cultured endothelial cells exposed to hypoxia and NG or HG conditions. Either under NG and HG exposure, 1h stimulation with Pyr-apelin-13 similarly raised AMPK*α*1/2 (Fig. 6A) and eNOS phosphorylation at Ser1177 (Fig. 6B) and Ser633 (Fig. 6C). In addition, eNOS mRNA expression in endothelial cells was increased by 2.6-fold (*P*=0.0008) and 1.8-fold (*P*=0.0410) following 24h stimulation of apelin under NG and HG conditions, respectively (Fig. 6D). As for the *in vivo* results, we observed a very similar pattern for RhoA/ROCK2 signaling pathway, apelin treatment increased ROCK-2 phosphorylation (Fig. 6E) and total protein expression (Fig. 6F) as well as RhoA mRNA expression (Fig. 6G) in both NG and HG conditions. These data demonstrate that apelin stimulation in endothelial cells, through APJ receptor, promotes signaling pathways that are not affected by hyperglycemia.

**Figure 6.**
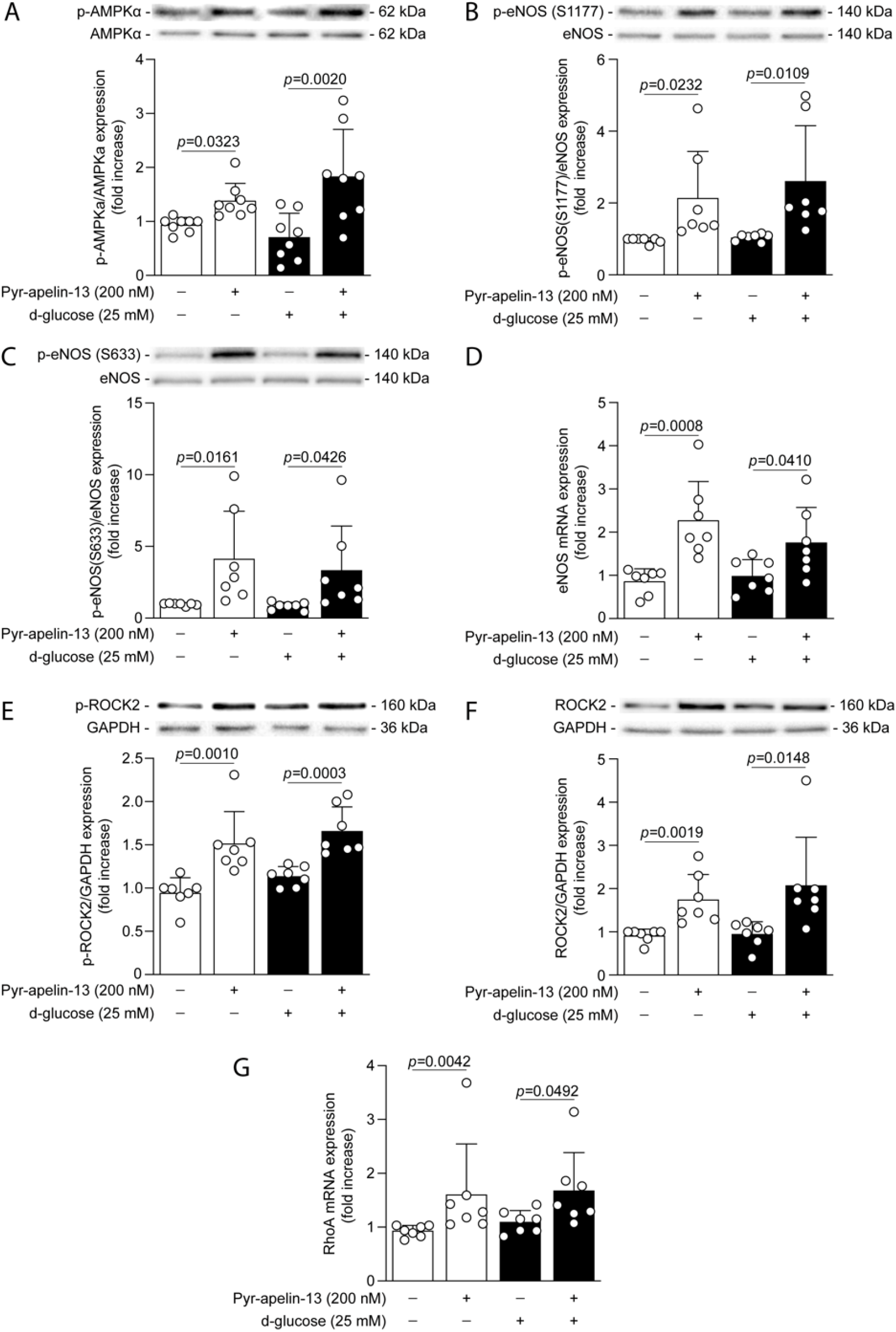
Pyr-apelin-13 enhanced the signaling pathways involved in cytoskeleton dynamics and proangiogenic actions in cultured endothelial cells exposed to high glucose conditions and hypoxia. BAECs were exposed to normal glucose (NG; 5.6 mmol/L; white bars) or high glucose (HG; 25 mmol/L; black bars) concentrations for 48h and then stimulated with Pyr-apelin-13 for 1h **(A-C, E** and **F)** or 24h **(D** and **G)**. BAECs were exposed to hypoxia (1% O_2_) for the last16h of treatment. Protein expression of **(A)** phospho-AMPK*α*1/2, **(B)** phospho-eNOS at Ser1177, **(C)** phospho-eNOS at Ser633, **(E)** phospho-ROCK-2 and **(F)** total protein expression of ROCK2 was detected by immunoblot analysis and the densitometry quantification was measured. mRNA levels of **(D)** eNOS and **(G)** RhoA. GAPDH gene was used for mRNA normalization. Results are presented as the mean ± SD of 8 (**A**) and 7 (**B**-**G**) independent cell experiments.

## Discussion

Patients with diabetes and PAD are 5 to 15 times more likely to undergo major amputation due to vascular abnormalities resulting from diabetes^37^. Indeed, diabetes impairs the angiogenic response, leading to impaired wound healing and collateral vessel formation^38^. Endogenous apelin has been shown to participate in the collateral vessel formation processes during ischemia^21^. Previous studies reported that apelin treatment or apelin gene therapy improved limb reperfusion after femoral artery ligation in nondiabetic animal models^22,23^. However, very few studies have characterized the impact of the apelinergic system on the angiogenic response under diabetic conditions. Our study demonstrated for the first time that apelin delivery significantly improved blood flow reperfusion and functional recovery of the hindlimb following ischemia in a diabetic condition.

The vascular endothelial growth factor (VEGF) is one of the most extensively studied growth factors in relation to physiological angiogenesis. VEGF stimulates endothelial cell proliferation, migration, differentiation and survival, leading to new blood vessel formation. Decreased mRNA and protein expression of VEGF-A/VEGFR-2 have been described in the muscle of diabetic mice and is associated with impaired revascularization following ischemia^7,26^. Although preclinical studies using VEGF therapies in animal models of hindlimb ischemia were promising^39^, all phase II clinical trials failed to significantly reduce the amputation rate in patients suffering of PAD^40^. A possible reason for these clinical disappointments may be caused by an impairment in VEGF receptors and downstream signaling (Akt, eNOS) activation in response to ligand binding created by high glucose exposure, a phenomenon highlighted in several preclinical studies^7^. Our study corroborated these findings that hyperglycemia prevented the activation and phosphorylation of VEGFR-2 downstream effector Akt, the proliferation, migration and tubule formation in response to VEGF stimulation in endothelial cells. These results underline a resistance phenomenon to VEGF actions induced by diabetes, preventing the natural mechanism of angiogenesis in response to tissue ischemia. In our study, we have recreated the ischemic state found in PAD and observed that Pyr-apelin-13 stimulation in endothelial cells exposed to the combination of hypoxia and NG or HG concentrations enhanced Akt and eNOS phosphorylation, promoted cell proliferation, migration and tubule formation. Moreover, our results indicated that Pyr-apelin-13 delivery in diabetic mice markedly increased Akt and eNOS phosphorylation, as well as eNOS protein expression in ischemic muscle. VEGFR2 (Flk-1) and PDGFR-*β* are not direct targets of APJ/apelin signaling pathways, but interestingly, we identified that apelin administration increased VEGF-A, VEGFR-2, PDGF-B and PDGFR-*β* mRNA expression in diabetic ischemic muscle. Taken together, our data suggest that Pyr-apelin-13 delivery circumvents the resistance mechanism induced by hyperglycemia, providing an effective angiogenic response under hypoxic conditions.

The apelinergic system’s implication in angiogenesis has been demonstrated by the presence of apelin/APJ in tip cells and stalk cells, which are responsible for guiding the new vessel formation toward the gradient of proangiogenic factors and for the elongation of the sprouting vessel^41^. Furthermore, other studies reported apelin/APJ upregulated expression under hypoxic and normoglycemic conditions in cultured endothelial cells, smooth muscle cells and peripheral mononuclear cells, as well as in mice lungs following ischemia^42^. The presence of hypoxia response element (HRE) sequence was found on apelin and APJ genes, making them targets of hypoxia-inducible factor 1 (HIF-1), a transcription factor regulating gene expression in response to hypoxia. Few studies investigated the basal level of expression of the apelinergic system in nonischemic tissues from diabetic animals and reported a decreased expression of apelin and/or APJ in the heart, skeletal muscle and serum^27,29^. Surprisingly, despite the presence of tissue ischemia, our results demonstrated that diabetes prevents the normal expression of APJ/apelin in the adductor muscle. Interestingly, we showed that administration of Pyr-apelin-13 in diabetic animals enhanced APJ and apelin expression, improving endothelial function and angiogenesis. Collectively, these data suggest that the repressive effect of diabetes on the apelinergic system expression can be overcome by apelin administration, resulting in a proper recovery from tissue ischemia.

AMPK is a metabolite-sensing protein kinase mostly known for its metabolic effects, such as increasing glucose uptake, fatty acid oxidation and inhibition of lipogenesis, but several studies emphasized its role during angiogenesis. Indeed, Nagata and al. reported that AMPK activation is essential for angiogenesis under hypoxic conditions, but dispensable to the angiogenic response under normoxic conditions^43^. As Akt, AMPK mediates the activity of eNOS by phosphorylation at Ser1177, which characterized its activation state, to induce NO release, promoting vasodilatation and proangiogenic actions^32^. Interestingly, AMPK but not Akt is able to phosphorylate eNOS in Ser633, a phosphorylation site reported to be an effective indicator of eNOS-induced NO bioavailability, resulting in enhanced endothelial cell migration and tubule formation^33^. Apelin has been reported to induce AMPK, Akt and eNOS phosphorylation under normal glucose concentrations^44^. Here, we demonstrated that these effects were preserved in hypoxic endothelial cells and ischemic muscle following apelin stimulation despite being exposed to HG levels and diabetes. Taken together, our results demonstrated that apelin treatment, under hypoxic and HG or diabetic conditions, induced eNOS activation and proangiogenic actions through two different phosphorylation sites involving Akt and AMPK activation.

Endothelial cell mobility involves the sensing of proangiogenic signals by filopodia, the formation, protrusion and extension of cytoskeleton projections, the assembly/disassembly of focal adhesions, the contraction of the cell body enabling forward movement and rear release. All these processes are mediated by the Rho small GTPase family^45^, a classical pathway downstream of the small G*α*_12/13_ protein recruitment following GPCRs activation^35^. More specifically, RhoA and its protein kinase effector ROCK regulate actin-regulating proteins, allowing anchoring of stress fibers to the membrane, and mediate the formation and contraction of the stress fibers and rear pulling during migration^45^. ROCK protein kinases are necessary for normal angiogenesis during embryonic development since ROCK1 or ROCK2 knockout mice died in utero due to impaired vasculature development^46^. Furthermore, inhibition of RhoA/ROCK signaling pathway suppresses VEGF-induced endothelial cell migration and tube formation, and angiogenesis *in vivo*^*47*^. Compared to ROCK1, ROCK2 is preferentially expressed in vascular cells^48^, and gene knockdown of ROCK2, but not ROCK1, reduced endothelial cell tube formation *in vitro* and vascular density in mice lungs^47^. However, no study demonstrated concrete evidence of the activation of this signaling pathway following apelin stimulation. To our knowledge, we demonstrated for the first time that apelin stimulation, through APJ receptor, led to the activation of the RhoA/ROCK signaling pathway in the ischemic muscle from diabetic mice and in endothelial cells exposed to HG levels. These results highlight the role of the apelinergic system on the actin cytoskeleton remodeling in endothelial cells, facilitating angiogenesis following ischemia.

Pain in the legs is one of the most common symptoms of PAD when exercising, which reduces walking distance. Our results clearly demonstrated the association between poor blood flow and reduced walking distance since diabetic mice displayed a 39% lower blood flow recovery and a 91% decreased running distance compared to nondiabetic controls. The increased blood flow reperfusion we observed following apelin treatment was supported by increased capillary density in the ischemic muscle compared to untreated diabetic mice. This is the first time that, in addition to improving blood flow reperfusion, sustained apelin delivery resulted in enhanced motor function of the ischemic limb in diabetic mice. Despite the improved functional recovery of the limb with apelin, an important gap remains compared to nondiabetic mice. One hypothesis could be nerve degeneration caused by diabetes, leading to typical symptoms of peripheral neuropathy such as muscle weakness, extreme sensitivity, and numbness. Interestingly, a recent study reported apelin’s neuroprotective and regenerative effects in a nondiabetic focal cerebral ischemia mice model^49^. However, further investigations will be required to evaluate the potential effects of apelin in diabetic neuropathy.

In conclusion, we demonstrated that, unlike VEGFR signaling pathways, the apelin/APJ pathways are not affected by hyperglycemia, making the apelinergic system a potential target for angiogenic therapy. Furthermore, we demonstrated the therapeutic effect of Pyr-apelin-13 in diabetic PAD, since its prolonged administration resulted in improved blood flow reperfusion, vascular density and motor function in diabetic mice following hindlimb ischemia. However, apelin’s short plasma half-life may represent a challenge for its use as a pharmacological treatment. Thus, research must continue to develop more stable apelinergic agonists to improve the long-term angiogenic response in diabetes^50^.

## Nonstandard Abbreviations and Acronyms

Akt: Protein kinase B
AMPK: AMP-activated protein kinase
APJ: Apelin receptor
BAEC: Bovine aortic endothelial cell
eNOS: Endothelial nitric oxide synthase
GPCR: G protein-coupled receptors
HG: High glucose
HIF-1*α*: Hypoxia-inducible factor-1*α*
NG: Normal glucose
NO: nitric oxide
PAD: Peripheral artery disease
PDGF: Platelet-derived growth factor
PDGFR: Platelet-derived growth factor receptor
Pyr-apelin-13: Pyroglutamyl-apelin-13
RhoA: Ras homolog family member A
ROCK: Rho-associated coiled-coil kinase
VEGF: Vascular endothelial growth factor
VEGFR: Vascular endothelial growth factor receptor

## Acknowledgments

S.R., T.B. and E.B. performed experiments. K.T., P-L.B. and É.M. synthesized and generously provided Pyr-apelin-13 peptide for the experiments. S.R., F.L. and P.G. analyzed the data and wrote the manuscript. P-L.B. and M.A-M. reviewed the manuscript and provided comments. The authors gratefully acknowledge and dedicate this article to Prof. Éric Marsault, who passed away^51^. Marilène Paquette (Histology Core, University of Sherbrooke) is gratefully acknowledged for her assistance with the histology technics. All authors approved the submitted manuscript and support its conclusion. P.G. is the guarantor of this work, had full access to all the data and takes full responsibility for the integrity of the data and the accuracy of data analysis.

